# Co-delivery of Paclitaxel and Cannabidiol in Lipid Nanoparticles Enhances Cytotoxicity Against Melanoma Cells

**DOI:** 10.1101/2024.10.15.618512

**Authors:** Fabíola V. de Carvalho, Gabriela Geronimo, Ludmilla D. de Moura, Talita C. Mendonça, Márcia Cristina Breitkreitz, Eneida de Paula, Gustavo H. Rodrigues da Silva

## Abstract

Although chemotherapy regimens are well-established in clinical practice, chemoresistance and adverse side effects pose significant obstacles in cancer treatment. Paclitaxel (PTX), a widely used chemotherapeutic agent, faces formulation challenges due to its poor solubility and permeability. Research suggests that the phytochemical Cannabidiol (CBD) holds potential not only in targeting cancer cells but also in alleviating pain and nausea, thereby improving the quality of life for cancer patients. However, CBD’s clinical application is also limited by its poor solubility, low bioavailability, and susceptibility to oxidation. Nanostructured lipid carriers (NLCs) represent a promising drug delivery system for hydrophobic compounds like PTX and CBD and allows their co-encapsulation. Nonetheless, achieving a stable formulation requires identifying suitable preparation methods and excipients. The aim of this study was to develop and optimize an NLC formulation for the co-encapsulation of PTX and CBD. Using factorial design, an optimized formulation was obtained with homogeneous particle sizes (200 nm), negative zeta potentials (−17 mV), particle concentration of 10^13^ particles/mL, spherical morphology (TEM images), and a lipid core with low crystallinity (as confirmed by XRD). To evaluate the therapeutic potential of the drug combination, cell viability assays were conducted on murine melanoma cells (B16-F10) at different exposure times (24 and 48 hours). The NLC-CBD-PTX formulation significantly reduced cell viability in a time- and concentration-dependent manner, demonstrating at least 75% greater activity at 24 hours compared to each drug individually whether free (PTX, CBD) or encapsulated (NLC-PTX, NLC-CBD). This indicates a synergistic effect of the encapsulated drugs on cytotoxicity. In conclusion, an innovative pharmaceutical formulation co-encapsulating PTX and CBD was validated, showing potential to enhance antitumor efficacy, overcome chemoresistance, reduce side effects, and broaden therapeutic applications. The resulting NLCs exhibited favorable physicochemical properties, supporting their suitability for various routes of administration.

## 1. INTRODUCTION

According to the World Health Organization, approximately 19 million people were diagnosed with cancer in 2020, resulting in around 10 million deaths [1]. In the United States, estimates for 2024 indicate 2,001,140 new cancer cases and 611,720 deaths related to the disease [2]. Globally, it is projected that by 2030, there will be over 25 million new cancer diagnoses [3]. In Brazil, the occurrence of 704,000 new cancer cases is estimated for the period from 2023 to 2025 [3]. These statistics underscore the importance of ongoing efforts in research, prevention, and treatment to combat cancer.

Chemotherapy treatment protocols are well established in clinical practice. Paclitaxel (PTX) is a broad-spectrum chemotherapeutic agent approved by the Food and Drug Administration (FDA) in 1992 for the treatment of various types of cancer, from early to advanced stages [4,5]. Derived from the plant *Taxus brevifolia*, PTX belongs to taxane class antineoplastics and exhibits cytotoxic activity against many solid tumors [6]. By binding to the N-terminal amino acids of the tubulin β-subunit, PTX stabilizes the main protein of microtubules and enhances its polymerization, disrupting mitosis in the G2/M phase and leading to cell death [6,7]. However, being a class IV drug in the Biopharmaceutics Classification System, PTX has low aqueous solubility and permeability [4,6,8–10]. Commercial pharmaceutical formulations, such as TAXOL^®^, contain non-inert excipients (ethanol and polyoxyethylated castor oil) that increase PTX toxicity, resulting in severe side effects such as cardiovascular issues, neutropenia, and central or peripheral neurotoxicity, limiting its clinical applicability [11–13]. In the context of chemotherapy, encapsulation in drug delivery systems (DDS) can minimize these undesirable effects by providing sustained drug release to avoid plasma spikes and targeting the drug to tumor cells [14].

Cannabidiol (CBD) has been extensively studied for the treatment of various conditions and diseases such as chronic pain, inflammation, and cancer [15–21]. A recent review highlighted the effects of CBD on human cancer cells from various tissues, including gastrointestinal, genital, mammary, respiratory, nervous, hematopoietic, and skeletal systems, showing a decrease in cell viability, proliferation, migration, inflammation, and metastasis [15]. In the case of melanoma, recent studies have demonstrated the potential antineoplastic effects of CBD through various molecular mechanisms, primarily involving reduction of tumor cell viability, proliferation, migration, and angiogenesis, as well as induction of apoptosis in preclinical models [22– 24]. Despite the numerous pharmaceutical activities attributed to CBD, its clinical applicability is hindered by low aqueous solubility (0.1 µg/mL), limited bioavailability (6%), and susceptibility to oxidation [25,26], demanding the development of strategies to enhance its clinical use [27].

Drug delivery systems (DDS) have gained increasing importance in the research and development of new medications due to their numerous advantages. These include protection of the drug from enzymatic degradation, prolonged circulation time in the bloodstream, reduced therapeutic doses, ease of administration, and lower toxicity with increased bioavailability. Additionally, their small size allows for greater accumulation in target tissues, enhancing therapeutic efficacy [28]. Nanostructured lipid carriers (NLC) are lipid nanocarriers composed by a lipid core (mixture of solid and liquid lipids at room/body temperature) stabilized by a surfactant. The lipid blend makes the core less ordered, allowing for a higher drug upload and minimizing drug expulsion during storage [29,30].

Chemoresistance poses a significant challenge in the effective treatment of cancer, as tumor cells develop resistance to chemotherapy drugs [31,32]. DDS may overcome drug resistance, enhancing drug delivery and improving treatment efficacy [22,33]. Indeed, when it comes to encapsulating chemotherapy drugs in NLCs, they have shown to surpass the limitations of traditional chemotherapy by enhancing drug loading capacity, stability, targeted delivery and chemoresistance [34,35]. Additionally, NLCs can be tailored to co-encapsulate multiple therapeutic agents, such as chemotherapeutic drugs and genetic material or metabolic modulators or siRNA, to achieve synergistic effects and combat drug resistance [36,37].

Despite the scientific advancements in the field, the development of more efficient antitumor formulations that are selective to the target cell, water-soluble for parenteral application, and with low systemic toxicity remains a significant challenge. Therefore, the aim of this study was to combine PTX and CBD in NLC to attain enhanced anticancer properties and reduce side effects, such as PTX-induced peripheral neuropathy. Additionally, cannabinoids may contribute to a better quality of life for cancer patients, offering palliative effects such as pain relief and nausea reduction [38]. In this study, we developed an optimized NLC-CBD-PTX formulation using experimental design, characterized it through techniques such as DLS, NTA, TEM, and XRD, and evaluated its stability. Finally, we assessed the cytotoxicity of the co-encapsulated actives in melanoma cells to determine their efficacy in anti-tumor activity.

## 2. MATERIALS AND METHODS

### 2.1 Materials

Paclitaxel powder (PTX) was a gift from Cristália Prod. Quim. Farm. Ltda (São Paulo, SP, Brazil) and myristyl myristate (MM) was donated by Croda do Brasil Ltda (Campinas, SP, Brazil). Commercial PTX (TAXOL^®^) was purchased in Brazilian market. Cannabidiol oil was purchased from Prati-Donaduzzi & Cia Ltda (Toledo, PR, Brazil). Soy L-α-Phosphatidylcholine (SPC) was from Avanti^®^ Polar Lipids, Inc. (Alabaster, AL, USA). Pluronic F-68 (P68), Dulbecco’s Modified Eagle Medium (DMEM), fetal bovine serum (FBS), 3-(4,5-dimethylthiazol-2-yl)-2,5-diphenyltetrazolium bromide (MTT), penicillin, streptomycin sulphate and trypsin were supplied by Sigma Chem. Co. (St. Louis, MO, USA). Dimethyl sulfoxide (DMSO) was purchased from Laborclin (Pinhais, PR, Brazil) and HPLC-grade methanol from J.T. Baker (Allentown, PA, USA). Melanoma cells (B16-F10 lineage) were purchased from American type culture collection (ATCC, Manassas, VA, USA). Deionized water (18 MΩ) was obtained with an Elga USF Maxima ultra-pure water purifier.

### 2.2. Methods

#### 2.2.1 NLC preparation

NLCs were prepared using a modified emulsification-ultrasonication technique [39]. Initially, PTX was dissolved in the lipids at a temperature 10 °C above the solid lipid’s melting point, with the addition of ethanol, and subjected to 10 minutes of heating and mechanical agitation in a water bath. Simultaneously, a surfactant solution was heated to the same temperature and both phases were mixed at high speed (10,000 rpm) for 3 minutes using an Ultra-Turrax blender (IKA Werke, Staufen, Germany). Subsequently, the mixture underwent 16 minutes of sonication in a Vibracell tip sonicator (Sonics & Materials Inc., Danbury, USA) operating at 500 W and 20 kHz, in alternating 30-second cycles. The resulting nanoemulsion was cooled to room temperature, to form the NLC.

#### 2.2.2. Composition optimization by experimental design

The selection of excipient quantities in this specific nanoparticle was investigated through factorial design using Design Expert® software (version 13, Stat-Ease, Inc., Minneapolis, USA). A full factorial design 2^3^ was developed with triplicates at the central point. Three independent variables were evaluated at two levels (high and low). The maximum and minimum factor-levels were determined through preliminary screening tests, taking into consideration previous design experiences of group. Table 1 displays the design conditions, with fixed quantities of 1g of cannabidiol oil (200 mg/mL) and 60 mg of PTX. The criteria for optimizing included particles of minimum sizes (versatility of application), minimal PDI (monodisperse systems), and higher ZP values, in module [40,41].

**Table 1.** Design of experiments. Experimental variables, levels, properties of interest (responses) and criteria for optimizing. Average diameters (size), polydispersity index (PDI) and Zeta potential (ZP).

| Experimental variables | Symbols in the program | Low level | High level |
| --- | --- | --- | --- |
| Myristyl Myristate (%) | A | 8 | 12 |
| L-α-Phosphatidylcholine (g) | B | 0.1 | 0.3 |
| Pluronic F68 (%) | C | 6 | 10 |
| Properties of interest |  | Criteria for optimization |  |
| Size (nm) |  | Minimum |  |
| PDI |  | Minimum |  |
| ZP (mV) |  | Maximum |  |

#### 2.2.3. Physicochemical characterization of NLC

##### 2.2.3.1. Measurement of particle size and polydispersity index

The average diameter and PDI of the NLCs were measured by dynamic light scattering (DLS). Dilutions of the different suspensions in deionized water (1000x) were measured in triplicate using the ZetaSizer Nano ZS90 equipment (Malvern Instruments, Malvern, Worcestershire, UK) at 25°C, coupled to a data acquisition system.

##### 2.2.3.2. Zeta Potential Measurement

ZP values were determined by Laser Doppler Microelectrophoresis using the ZetaSizer Nano ZS90 equipment (Malvern Instruments, UK). Measurements were performed in triplicate in appropriate polystyrene cuvettes, diluting the NLC suspensions in deionized water (1000x), at a temperature of 25°C [40]. The data were expressed as mean ± SD.

##### 2.2.3.3. HPLC method and determination of encapsulation efficiency

The drugs PTX and CBD were quantified by High-Performance Liquid Chromatography (HPLC) under the conditions described in Table 2. The equipment used was a Waters Breeze 2 high-performance liquid chromatograph (Waters Technol., MA, USA). The retention times for PTX and CBD were 4.9 and 20 minutes, respectively, (as shown in the chromatogram of Figure S1).

**Table 2.** Chromatographic conditions for the simultaneous quantification of PTX and CBD in NLCs.

| Parameters | Conditions |
| --- | --- |
| Column | C18 Gemini-NX 5μ 450 x 4.60 mm |
| Oven temperature | 30°C |
| Mobile phase | Acetonitrile:0.1% acetic acid, 70:30 (v/v) |
| Flow | 1 mL.min<sup>-1</sup> |
| Injection volume | 10 μL |
| Wavelength | 230 nm |

The percentage of encapsulation efficiency (%EE) was determined by the ultrafiltration-centrifugation method, using cellulose filters (10 kDa, Millipore) [40,42]. For this purpose, an aliquot of the formulation was added to the filtration unit coupled to Eppendorf tubes and centrifuged for approximately 20 minutes at 4100g. The amount of free antineoplastic in the filtrate was quantified by HPLC and the percentage of encapsulated antineoplastic was calculated according to Equation 1.

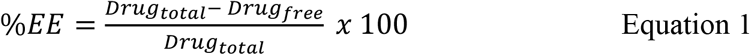

where Drug_total_ is the total amount of PTX or CBD quantified in the NLC suspension and Drug_free_ corresponds to the amounts of PTX or CBD in the filtrate.

##### 2.2.3.4. Nanoparticle tracking analysis

Nanoparticle Tracking Analysis (NTA) was used to determine the size, distribution and concentration of nanoparticles in the formulation, on an NS300 instrument (NanoSight, Amesbury, UK) equipped with a 532 nm laser. Based on the tracking of individual Brownian motion of nanoparticles, NTA is the only real-time technique that provides the nanoparticle concentration (number of particle/mL) [43]. Samples were diluted in deionized water (50,000x) and introduced into the sample holder using a syringe until it was filled completely. The measurements were performed at room temperature, in triplicate, with the data expressed as the mean ± SD.

The polydispersity or particle size distribution data were obtained by calculating the SPAN index, according to Equation 2 [44].

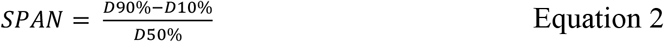

Where D10, D50 and D90 refer to the mean size of 10%, 50% and 90% of the particle population, respectively.

##### 2.2.3.5. X-Ray diffraction

The X-Ray diffraction (XRD) measurements were carried out using the D2-Phaser diffractometer (Bruker, Germany) under the following experimental conditions: temperature of 293 K, Cu-Kα radiation (λ=1.5418 Å) generated at 30 kV and 10 mA, continuous scan mode with a step time of 0.2 s, angular range (2θ) of 5–50° with an increment of 0.02°. The NLC the samples were freeze-dried before the analysis. The generated data were processed using Origin software, version 8.2.

##### 2.2.3.6. Transmission Electron Microscopy

To analyze the morphology of the nanoparticles, a Tecnai G2 Spirit BioTWIN Transmission Electron Microscope (FEI Company, USA) with an accelerating voltage of 60 kV was used. Before staining, NLC samples were diluted 33 times in deionized water and placed on a formvar film covered with a copper grid. Then, the samples were stained with 2% uranyl acetate. After washing with deionized water and drying at room temperature (24 h), the grid replica was prepared in a sample holder and placed in the vacuum chamber of the instrument. The ImageJ^®^ software (version 1.53k, NIH, Bethesda, MD, USA) was used to edit the images and measure the particles size [45].

#### 2.2.4 Evaluation of formulation stability

The stability of the optimized formulation and its control, stored at room temperature, was evaluated for 2 months, as an aqueous dispersion. The analyzed parameters were: size (nm), PDI and PZ (mV), in addition to the visual inspection (appearance) of the formulations.

#### 2.2.5. Cell viability assay

##### 2.2.5.1 Cell culture

Cells of the B16-F10 cell lineage (murine melanoma cells, ATCC CRL 6475^®^) were cultured in DMEM (Dulbecco’s modified Eagle’s medium) supplemented with 10% FBS (fetal bovine serum) and 1% antibiotic (penicillin and streptomycin) and brewed in an incubator (Shel Lab – CO_2_ incubator, USA) at 37 °C with atmospheric humidity, containing 95% air and 5% CO_2_. The culture medium was changed every two days and from the subconfluence of more than 70% of the flask, cell subcultures were performed with the aid of trypsin.

##### 2.2.5.2 Cell viability assay

96-well microplates were used for plating - at a cell density of 2 × 10^4^ cells/well (24 h) or 1 × 10^4^ cells/well (48 h) - incubated at 37 ºC, with 5% CO_2_, for 24 hours for cell adhesion to occur. After this period, the medium was replaced with medium containing the respective treatment groups diluted, at 9 increasing concentrations (2 a 2×10^−8^ mM), and these treatments were performed for 24 and 48 h. After the incubation period, the supernatant was removed, the wells were washed with 5 mM PBS and viability was assessed by the soluble MTT (3-(4,5-dimethylthiazol-2-yl)-2,5-diphenyl tetrazolium bromide) reduction test: 0.5 mg/mL of MTT was added to the plate that was kept in the absence of light for 3 hours, at 37 ºC. After this period, the medium was carefully removed and DMSO was added to solubilize formazan crystals (produced by the degradation of MTT by the action of mitochondrial dehydrogenases of viable cells [46]. Finally, the plates were shaken for 10 minutes and the absorbance corresponding to each well was read in a ELx800-GEN5RC Elisa plate reader (Life Res. Co. London, England) at λ = 570 nm. The values were expressed as a percentage of MTT reduction in relation to the control, in which the cells were not exposed to the treatment.

## 3. RESULTS AND DISCUSSION

### 3.1 Composition optimization by experimental design

The initial step in formulating the NLCs involved identifying the optimal lipid blend for encapsulation of the active ingredients. The solid lipid myristyl myristate was selected based on previous research conducted by the group with another taxane drug [47]. Similarly, the non-ionic surfactant Pluronic F-68 was chosen. As for the liquid lipid, we picked a commercial cannabidiol oil prepared in corn oil. To enhance the system’s stability, we incorporated SPC as a co-surfactant, commonly used in NLC designed for antitumor agents [34]. A modification in the traditional preparation method of NLCs through ultra-homogenization and sonication was the approach taken for solubilizing paclitaxel in the lipid phase, with a small amount of ethanol.

Although the lipid composition was predetermined, we employed experimental design to investigate the influence of each excipient on the desired properties of the formulation, including particle physical properties such as size, PDI, and Zeta potential. Additionally, we analyzed how variations in these factors affected the final visual characteristics of the formulation (e.g., liquid formulation, highly viscous formulation such as a gel, or the formation of precipitates) after one week of preparation. As a result, nine different formulation compositions were studied, and the findings from the experimental design are presented in Table 3.

**Table 3.** Results obtained from the 2^3^-factorial design, showing the independent variables (MM, SPC and P68) and the dependent variables (size, PDI, ZP) for the NLC-CBD-PTX development.

|  | Factor 1 | Factor 2 | Factor 3 | Response 1 | Response 2 | Response 3 |
| --- | --- | --- | --- | --- | --- | --- |
| Formulation | A: MM (%) | B: SPC (g) | C: P68 (%) | Size (nm) | PDI - | ZP (mV) |
| 1 | 8 | 0.1 | 6 | 226.7 | 0.195 | -3.9 |
| 2 | 12 | 0.1 | 6 | 199.1 | 0.193 | -8.5 |
| 3 | 8 | 0.3 | 6 | 191.7 | 0.186 | -11.7 |
| 4 | 12 | 0.3 | 6 | 195.6 | 0.164 | -10.3 |
| 5 | 8 | 0.1 | 10 | 181.1 | 0.248 | -6.8 |
| 6 | 12 | 0.1 | 10 | 174.7 | 0.195 | -2.7 |
| 7 | 8 | 0.3 | 10 | 172.6 | 0.306 | -10.0 |
| 8 | 12 | 0.3 | 10 | 173.4 | 0.232 | -5.9 |
| 9 | 10 | 0.2 | 8 | 201.7 | 0.169 | -16.1 |
| 10 | 10 | 0.2 | 8 | 207.2 | 0.206 | -13.1 |
| 11 | 10 | 0.2 | 8 | 195.0 | 0.175 | -9.3 |

Regarding the influence of excipient variation on the physical properties of particles, the parameters size, PDI, and ZP values of the formulations were analyzed immediately after preparation. The particle sizes ranged from 172.6 to 226.7 nm. The mathematical model generated was significant and well-fitted (ANOVA, Table S1). As expected, SPC and P68 had negative effects on size (Figure 1A and Table 4), meaning that higher concentrations resulted in smaller sized NLCs. since surfactant and co-surfactants act to reduce the surface tension between the aqueous and lipid phases, leading to smaller particles [48].

**Figure 1.**
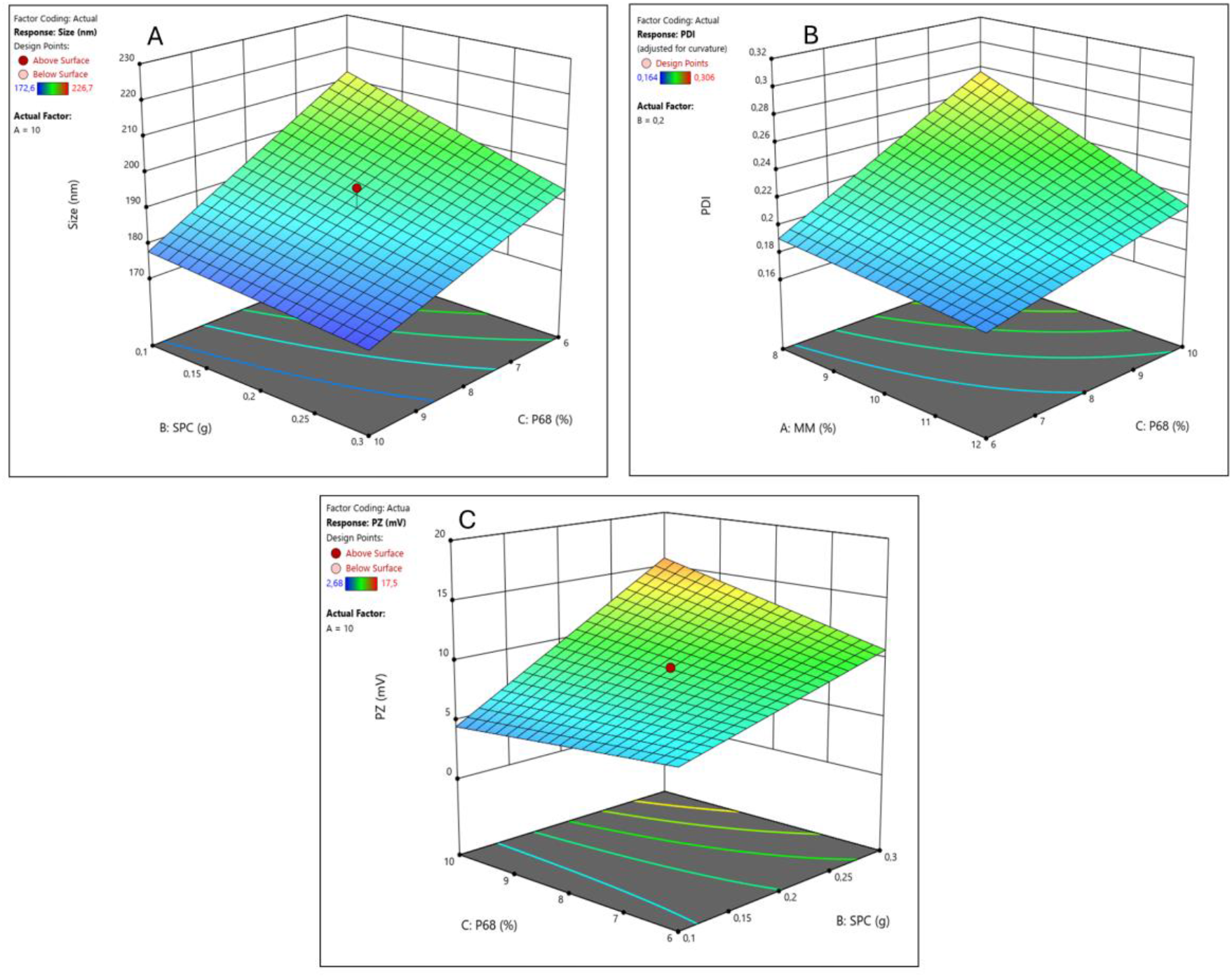
Factorial design results for the NLC-CBD-PTX system: response surfaces for size (**A**), PDI (**B**), and ZP (**C**).

**Table 4.** Components with significant effects, and their interactions, in each response analyzed for the NLC-CBD-PTX formulation. Factors: A= MM, B= SPC, C= P68.

| Response | Positive Effect | Negative Effect |
| --- | --- | --- |
| Size | - | B, C |
| PDI | C, BC | A, AC |
| ZP | B, BC, ABC |  |

PDI values ranged from 0.164 to 0.306, with four samples showing a PDI above 0.2 (samples 5, 7, 8, and 10), indicating polydisperse nanoparticles [42]. The mathematical model obtained was significant but required adjustment, with the curvature term added (see Table S2). Therefore, an increase in the response surface would be necessary to improve the mathematical model. The interaction between surfactant and co-surfactant had a positive effect on this response, while the interaction between solid lipid and surfactant had a negative effect, resulting in lower PDI values (see Figure 1B and Table 4).

The Zeta potential ranged from -2.7 to -16.1 mV. The mathematical model obtained was linear and showed no lack of fit (Table S3). Interestingly, the co-surfactant SPC alone and its interaction with other components had a positive effect on this response. In other words, the higher the amount of SPC, the higher the absolute ZP values (Figure 1C and Table 4). As a co-surfactant, this is a positive outcome as its addition to the system increased electrostatic repulsion between particles, potentially enhancing their colloidal stability. Moreover, in this case, surfactant P68 also played a crucial role in the steric stability of the formulation, essential for particle stability [49].

The expertise of the group indicates that some formulations may exhibit structural instability after one week of preparation, leading to phase separation, precipitation, or other visual phenomena. Therefore, the formulations prepared during the experimental design were monitored after one week to assess their visual stability. It was observed that all formulations prepared with a low level of P68 (6%) showed some form of visual instability after one week (Formulations 1 to 4, Table 3). According to the desirability graph of this experimental design (Figure S2), the optimized formulations (smaller size and PDI and higher Zeta potential [40,50]) should contain 6% of P68. However, low P68 concentration leads to long-term instability of the formulation. Due to this factor, we have selected the optimized formulations of the central point (8% P68) as they meet the desired criteria and remain visually stable. So, formulation number 9, belonging to the central point group, had its physical stability analyzed in terms of size, PDI, and ZP over a period of 60 days (Figure 2). Therefore, formulation 9 (with 10% MM, 0.2 g SPC, 1g CBD oil, 60 mg PTX and 8% P68) demonstrated stability for at least 60 days, with no significant changes observed in these parameters.

**Figure 2.**
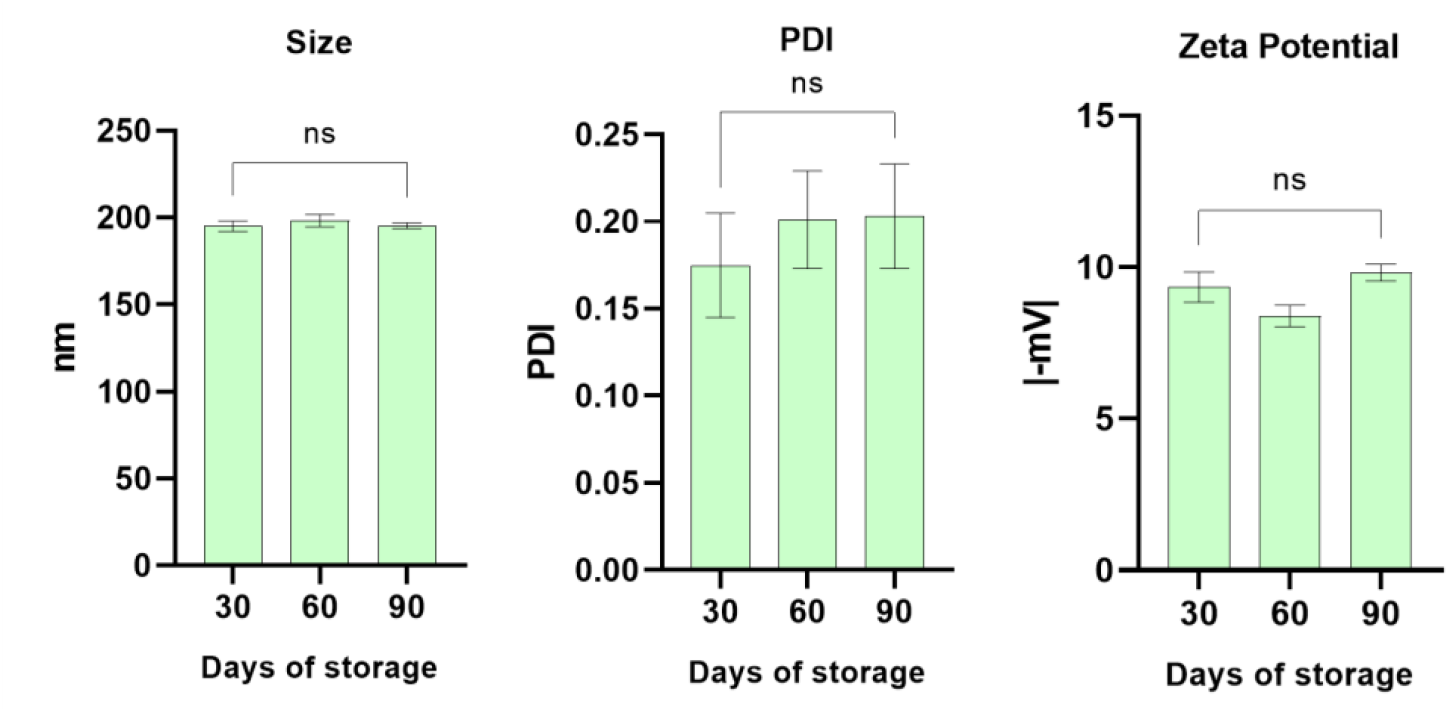
Physical stability in terms of size, PDI and ZP of the nanoparticles of the optimized NLC (formulation 9) containing cannabidiol and paclitaxel, under storage at room temperature for 60 days. ANOVA port hoc Tukey test: ns, non-significative.

### 3.2 Characterizations of optimized formulation

#### 3.2.1. DLS and entrapment efficiency

A fresh new batch of NLC formulations was prepared for the continuation of the study, based on the chosen composition determined by Design of Experiments: the optimized formulation for co-delivery of cannabidiol and paclitaxel (NLC-CBD-PTX) and control formulations, with each of the actives; paclitaxel (NLC-PTX) or cannabidiol (NLC-CBD). These formulations were characterized, and their physicochemical properties are given in Table 5. The NLC-CBD-PTX formulation had the largest size among all, reaching 206.5 nm and PDI values did not surpass 0.2, the limit to be considered a monodisperse population. ZP values were negative and only approach zero for NLC-PTX (−4 mV). Due to the hydrophobicity of the actives, the encapsulation efficiency was very high (≥ 98%) both for PTX and CBD, in the three formulations. Furthermore, co-encapsulation did not affect the %EE of the actives.

**Table 5.**
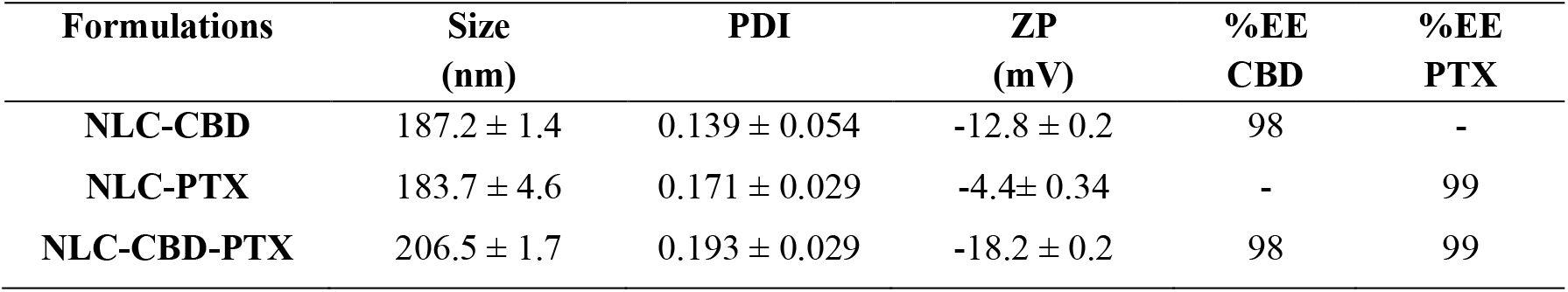
Physicochemical properties (size, PDI, ZP) and encapsulation efficiency (%EE) of PTX and CBD) in the formulation selected by factorial design (NLC-CBD-PTX) and its controls (NLC-CBD, NLC-PTX).

| Formulations | Size (nm) | PDI | ZP (mV) | %EE CBD | %EE PTX |
| --- | --- | --- | --- | --- | --- |
| NLC-CBD | 187.2 ± 1.4 | 0.139 ± 0.054 | -12.8 ± 0.2 | 98 | - |
| NLC-PTX | 183.7 ± 4.6 | 0.171 ± 0.029 | -4.4 ± 0.34 | - | 99 |
| NLC-CBD-PTX | 206.5 ± 1.7 | 0.193 ± 0.029 | -18.2 ± 0.2 | 98 | 99 |

#### 3.2.2. Nanoparticle tracking analysis

In the Nanoparticle Tracking Analysis (NTA) technique, application of the Stokes-Einstein equation allows for the determination of the size (nm) of each tracked particle. Furthermore, the specific and individual counting of particles in relation to volume enables a quick determination of nanoparticles concentration in each sample [51]. By analyzing the distribution of particles based on size (refer to Table 6), one can determine the uniformity of nanoparticles size, by the SPAN index that considers the D10, D50 and D90 the mean size of 10%, 50% and 90% of the nanoparticles, respectively accordingly to equation 2.

**Table 6.**
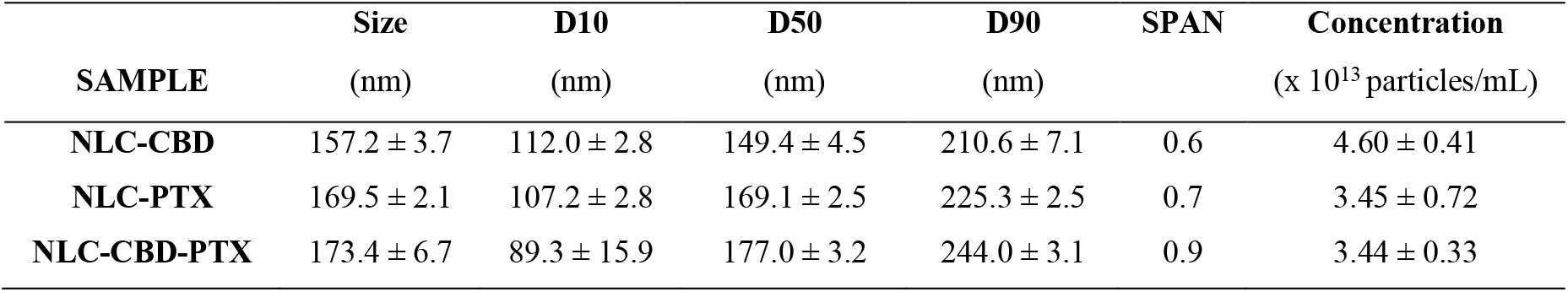
NTA Analysis: mean size and size dispersion at 10% (D10), 50% (D50), 90% (D90) of the NLC population, SPAN index (see text) and particle concentration in the optimized formulation and its controls (without paclitaxel or cannabidiol).

| SAMPLE | Size (nm) | D10 (nm) | D50 (nm) | D90 (nm) | SPAN | Concentration (x $10^{13}$ particles/mL) |
| --- | --- | --- | --- | --- | --- | --- |
| NLC-CBD | 157.2 ± 3.7 | 112.0 ± 2.8 | 149.4 ± 4.5 | 210.6 ± 7.1 | 0.6 | 4.60 ± 0.41 |
| NLC-PTX | 169.5 ± 2.1 | 107.2 ± 2.8 | 169.1 ± 2.5 | 225.3 ± 2.5 | 0.7 | 3.45 ± 0.72 |
| NLC-CBD-PTX | 173.4 ± 6.7 | 89.3 ± 15.9 | 177.0 ± 3.2 | 244.0 ± 3.1 | 0.9 | 3.44 ± 0.33 |

The mean diameters determined by NTA were slightly smaller but in good agreement with those obtained from DLS (Figure 2), as expected due to the differing principles of these techniques [42]. SPAN values below 1 indicate a uniform distribution of particle diameters [51,52]. Regarding the particle concentration, the formulations showed values on the order of 10^13^ particles/mL (Table 6). These concentrations are consistent with values reported in the literature, for similar NLC systems [40,53].

#### 3.2.3. X-Ray diffraction

The X-ray diffraction analysis (Figure 3) provided information into the crystallinity of the NLC lipid core [54], shedding light on the stability of the drug within the particles. Pure myristyl myristate, the key lipid excipient responsible for the solid core of the nanoparticles (28% in the dry mass), exhibited intense peaks at 21 and 24°, confirming its crystalline nature [55]. However, the intensity of these peaks decreased in the NLC formulations, indicating a reduction in the core’s crystallinity, probably because of the insertion of the actives in between MM, in the core of nanoparticles [56]. Similarly, the crystalline structure of P68, identified by peaks at 19 and 23°, was also diminished in the NLC formulations, suggesting its incorporation into the nanoparticle. Pure PTX displayed weak diffraction peaks at scattering 2θ angles at 5, 7, 10, and 13° [57], which were absent in the nanoparticles (NLC-PTX and NLC-CBD-PTX), indicating the partition of PTX into the NLC lipid matrix, considering its presence (16% of the NLC dry mass).

**Figure 3.**
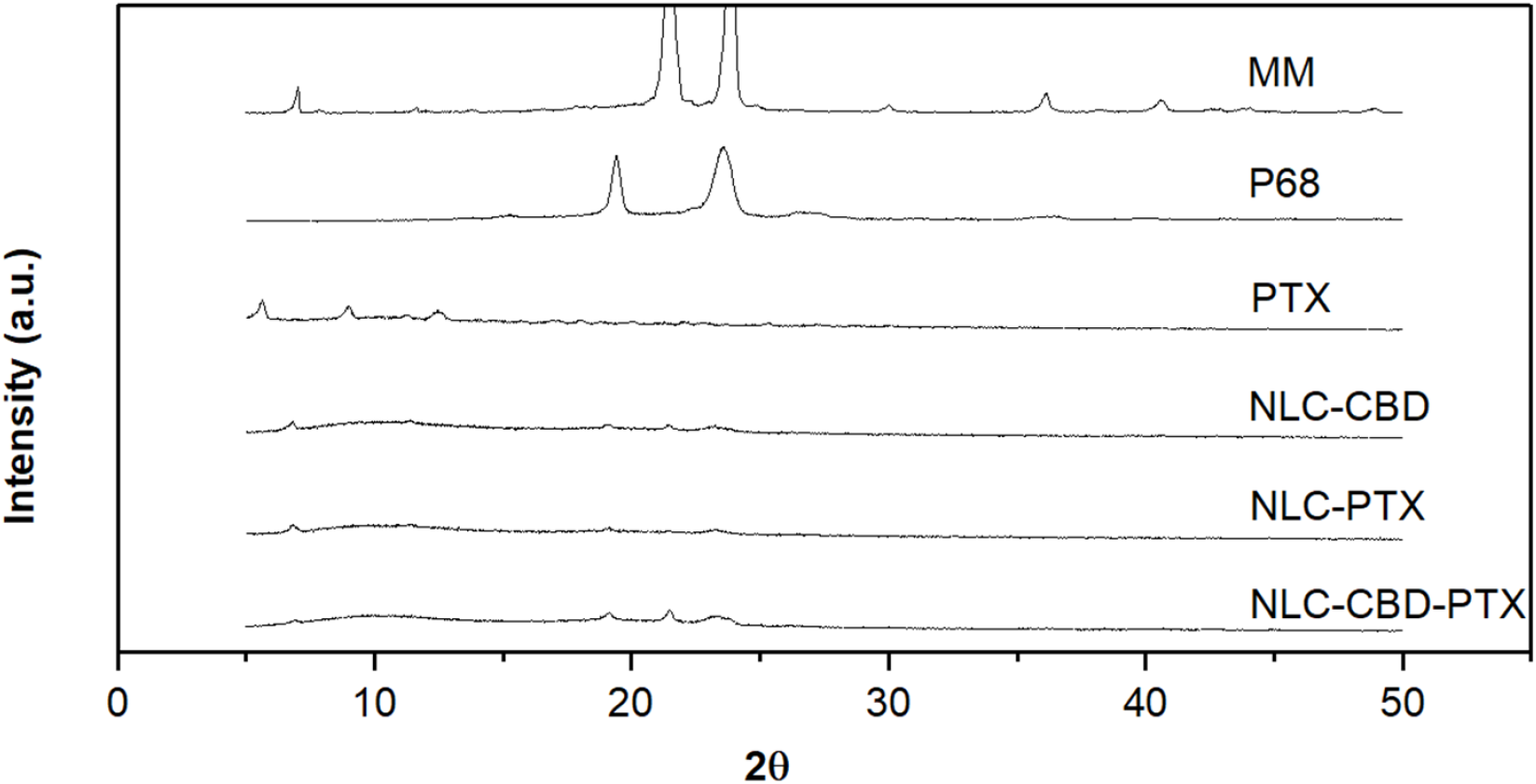
X-ray diffractograms of: pure myristyl myristate (MM), pure Pluronic F-68 (P68), pure paclitaxel (PTX), cannabidiol encapsulated in NLC (NLC-CBD), paclitaxel encapsulated in NLC (NLC-PTX) and the association of cannabidiol and paclitaxel in NLC (NLC-CBD-PTX). NLC formulations were freeze-dried before analysis. All diffractograms are in the same scale.

#### 3.2.4. Transmission Electron Microscopy

The Transmission Electron Microscopy (TEM) analyses provided valuable insights into the morphology of NLC and its nanometric size. Micrographies of diluted NLC formulations with and without PTX are shown in Figure 4, showing predominantly spherical nanoparticles of smooth surfaces, typical of lipid carriers. It is worth noting that addition of PTX did not change the morphology of the nanoparticles (Figures 4C and 4D). Their sizes, measured using the ImageJ^®^ software, were close to ∼200 nm, in good agreement with measurements obtained by DLS and NTA.

**Figure 4.**
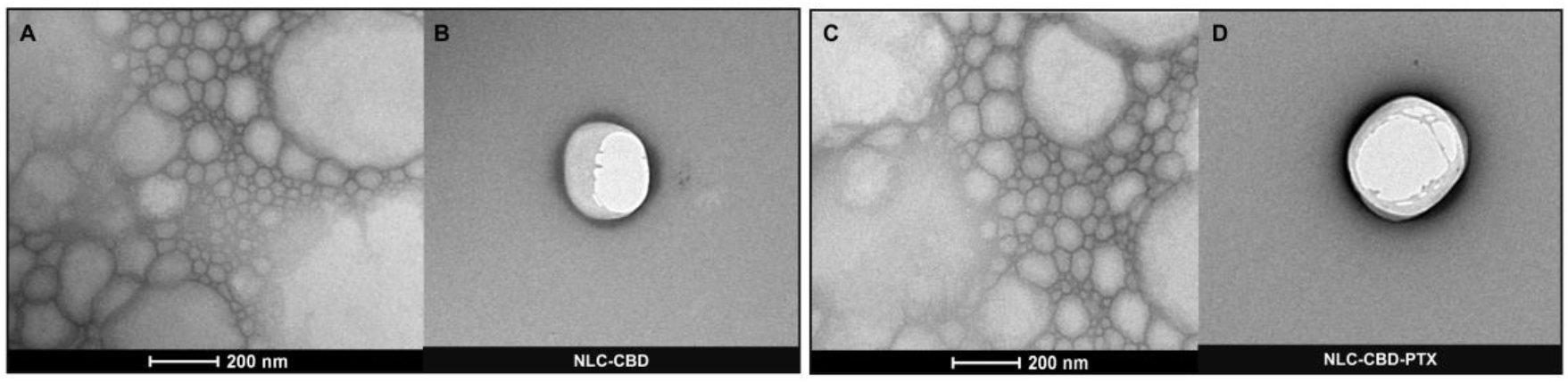
TEM micrographs of NLC formulations without PTX (NLC-CBD, **A, B**) and with PTX (NLC-CBD-PTX, **C, D**). Magnifications:18,500x (A e C) e 23,000x (B e D).

### 3.3. Cytotoxicity in melanoma cell line

To assess the impact of co-encapsulation on melanoma cells, the murine melanoma B16-F10 cell line was selected for its widespread use in *in vitro* testing and its ability to induce tumors *in vivo*. The cytotoxic effect was measured using the MTT assay [58,59], after 24 and 48 hours of nanoparticle exposure to the treatments.

Figure 5 A, B display the outcomes of treating B16-F10 cells for 24 h and 48 h, respectively, with formulations of free CBD oil and CBD oil encapsulated in NLC (NLC-CBD). This allowed us to observe the effect of CBD and its encapsulation on this cell line. Burch et al [60], demonstrated that CBD oil at a concentration of 0.4 mg/mL inhibits the *in vitro* growth of B16-F0 melanoma cells. In our study, using a more aggressive melanoma lineage, B16-F10, CBD oil was able to reduce viability by 50% (IC_50_) at ∼0.9 mg/mL after 24 h of exposure and ∼0.7 mg/mL after 48 h of exposure (Table 7). So, our results align with previous findings in the literature. Encapsulation of CBD in NLC increased its cytotoxicity (IC_50_ = 0.02 mg/mL after 24 h of exposure and 0.015 mg/mL after 48 h). A control NLC formulation, prepared without CBD, at the same excipients concentration and particle number, had a lesser effect on cytotoxicity compared to NLC-CBD, confirming the action of encapsulated CBD on the studied cell line (data not shown).

**Table 7.** Half-maximal inhibitory concentration (IC_50_) values determined for cannabidiol oil free (CBD) or encapsulated in NLC (NLC-CDB); commercial paclitaxel (TAXOL^®^) or PTX encapsulated in NLC (NLC-PTX) and the association of cannabidiol and paclitaxel in NLC (NLC-CBD-PTX) against B16F10 cells, after 24 and 48 h of treatments, as measured by the MTT assay. Analyses performed with GraphPad Prism 9.3.0 software with values of the curves in Figure 5.

| Formulation | $IC_{50}$ | | | | |
| --- | --- | --- | --- | --- | --- |
|  | CBD<br>(mg/mL) |  | PTX<br>(mM) |  |  |
|  | 24h | 48h | 24h | 48h |  |
| CBD | ~ 0.9 | ~ 0.7 | - | - |  |
| NLC-CBD | 0.02 ± 0.01 | 0.015 ± 0.003 | - | - | |
| TAXOL® | - | - | 4 × 10<sup>-3</sup> ± 7 × 10<sup>-3</sup> | 8 × 10<sup>-8</sup> ± 5 × 10<sup>-5</sup> | |
| NLC-PTX | - | - | 0.14 ± 0.16 | 4 × 10<sup>-3</sup> ± 3 × 10<sup>-3</sup> | |
| NLC-CBD-PTX | 0.005 ± 0.003 | 0.0002 ± 0.0003 | 8 × 10<sup>-3</sup> ± 4 × 10<sup>-3</sup> | 1 × 10<sup>-4</sup> ± 5 × 10<sup>-5</sup> | |

**Figure 5.**
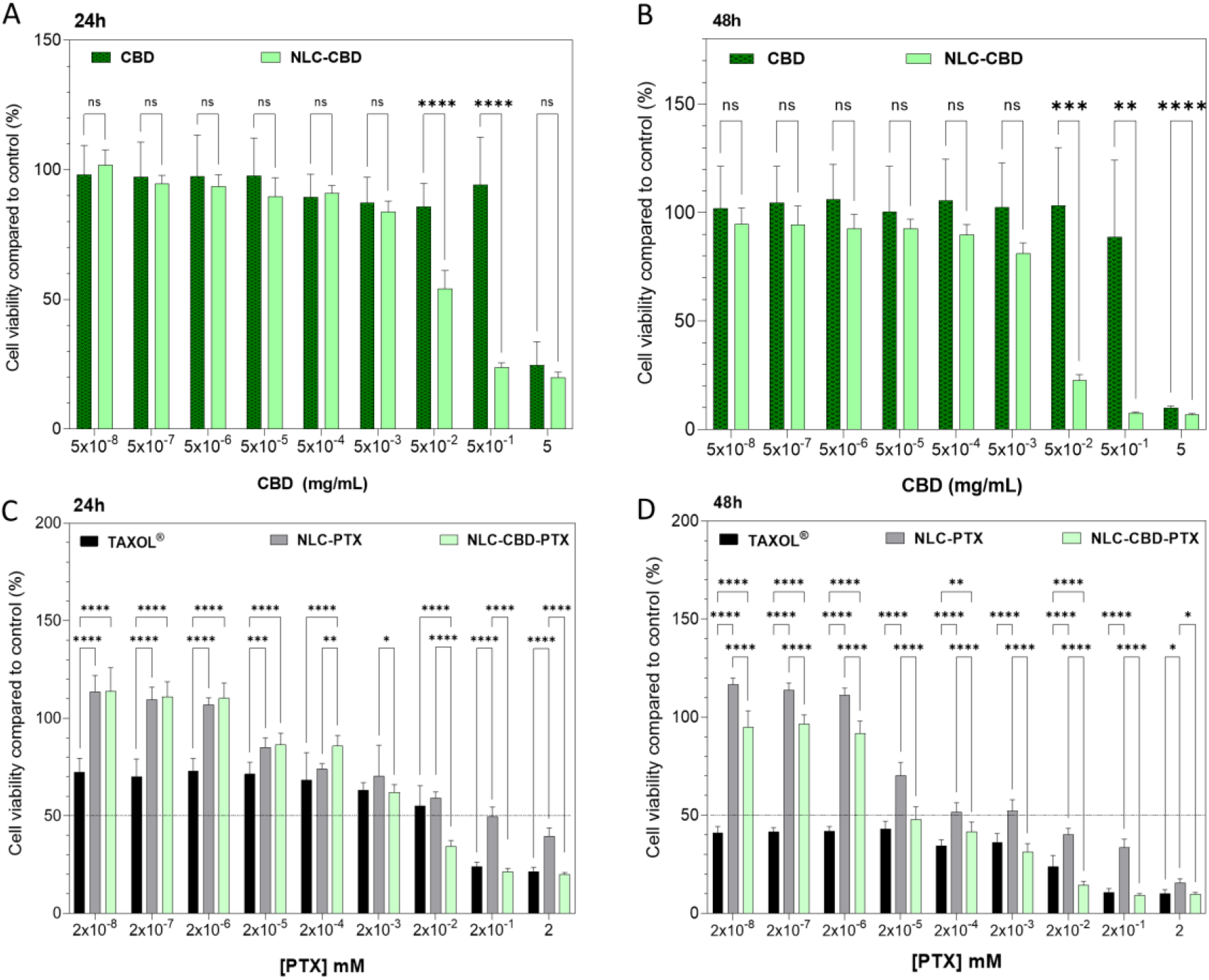
Cell viability (MTT assay) of melanoma strain (B16F10 cells) treated for 24 h (**A, C**) and 48 h (**B, D**) with cannabidiol oil (CBD) or encapsulated in NLC (NLC-CDB), paclitaxel commercial (TAXOL^®^) or encapsulated in NLC (NLC-PTX) and the association of cannabidiol and paclitaxel in NLC (NLC-CBD-PTX). Results expressed as mean ± SD (n = 12). Statistical analysis by Two-way ANOVA plus Tukey-Kramer post hoc. * p < 0.05; ** p < 0.01; *** p < 0.001, **** p < 0.0001.

The formulations containing free paclitaxel (TAXOL^®^) and encapsulated (NLC-PTX) were tested after 24 and 48 hours of exposure (Figure 5C and D, respectively). The reference drug TAXOL^®^ is a micellar formulation of Cremophor EL® (polyoxyethylated castor oil) in an ethanol:water solution, which evidently causes intense cell death even at low concentrations: IC_50_ of 4 × 10^−3^ and 8 × 10^−8^ mM at 24 and 48 hours, respectively. PTX encapsulated in NLC mitigates the drastic cell death seen in the TAXOL^®^ formulation, likely due to its less toxic excipients, showing an IC_50_ of 1.4 × 10^−1^ and 4.0 × 10^−3^ mM after 24 and 48 hours of exposure, respectively.

The optimized formulation (NLC-CBD-PTX) demonstrated higher cytotoxicity compared to formulations of NLC with the active ingredients (NLC-CBD and NLC-PTX) separated. There was a 75% decrease in the IC_50_ of CBD (0.005 mg/mL) after 24 hours of exposure and a 98% decrease (0.0002 mg/mL) after 48 hours of exposure. As for PTX, the decrease in IC_50_ was of 94% and 98% after 24 and 48 hours of exposure, respectively. In other words, a lower concentration of both active ingredients is required to cause 50% of cell death when they are co-encapsulated in the same formulation. Therefore, these data suggest that there may be a synergistic action between the PTX and CBD leading to cancer cell death.

## 4. CONCLUSION

Based on the obtained results, nanostructured lipid carriers co-encapsulating paclitaxel and cannabidiol were successfully developed and characterized, aiming to enhance the anticancer properties and reduce the side effects associated with chemotherapy. Physical-chemical analyses conducted using techniques such as DLS, NTA, TEM, and XRD, demonstrated the feasibility of the developed system. Cytotoxicity assays confirmed the antitumor efficacy and synergistic action of the co-encapsulated actives, highlighting the potential of this system for future *in vivo* testing, particularly in chemo resistant tumors. Additionally, the addition of CBD to conventional (PTX) treatment in a single formulation not only offers benefits in chemotherapy due to the need for a single administration but also presents a promising strategy to reduce PTX-associated side effects, such as peripheral neuropathy. Furthermore, this study demonstrates the therapeutic potential of cannabidiol when combined with a classic antineoplastic, offering not only a pharmaceutical option just in palliative care situations, but also the ability to act in earlier stages of chemotherapy treatment. So, these advancements contribute to the development of more effective and less aggressive treatments for oncology patients, underscoring the potential of NLCs as therapeutic vehicles.

## AUTHOR CONTRIBUTIONS

All authors discussed the results and commented on the manuscript. F.V.C. and G.G.: methodology, data curation, investigation, writing/original draft preparation. L.D.M.: methodology and investigation. T.C.M.: TEM analysis. E.P.: conceptualization, project administration, funding acquisition, resources and writing - review & editing. G.H.R.S.: conceptualization, supervision, methodology, data curation, visualization, resources, investigation and writing/original draft preparation.

## ACKNOWLEDGMENTS

The authors also thank Cristália Prod. Quím. Farm. Ltda for providing paclitaxel (powder) and Croda for donating myristyl myristate. We also thank the access to equipment and assistance provided by the Electron Microscope Laboratory – IB/Unicamp (LME/UNICAMP).

## DECLARATION OF AI-ASSISTED TECHNOLOGIES

During the preparation of this work the authors the authors exclusively utilized artificial intelligence in the writing phase to enhance the clarity and linguistic quality of the manuscript.

