## Supplementary Materials for "Co-delivery of Paclitaxel and Cannabidiol in Lipid Nanoparticles Enhances Cytotoxicity Against Melanoma Cells"

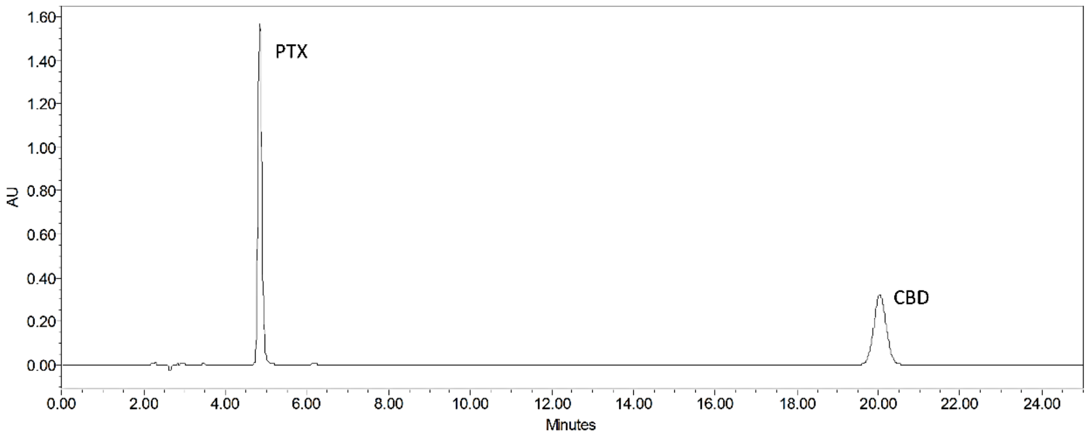


**Figure S1 -** Chromatogram showing the separation of paclitaxel (PTX) and cannabidiol (CBD) peaks at 4.9 and 20 min, respectively, in the NLC-CBD-PTX sample.

**Table S1**. ANOVA test for factorial model of the response: Size of nanoparticles.

| Source | Sum of Squares | df | Mean Square | F-value | p-value | |
|---|---|---|---|---|---|---|
| **Model** | 2352.32 | 7 | 336.05 | 17.46 | 0.0194 | significant |
| A-MM | 107.31 | 1 | 107.31 | 5.58 | 0.0993 | |
| B-SPC | 291.61 | 1 | 291.61 | 15.15 | 0.0301 | |
| C-P68 | 1548.46 | 1 | 1548.46 | 80.47 | 0.0029 | |
| AB | 187.21 | 1 | 187.21 | 9.73 | 0.0525 | |
| AC | 40.95 | 1 | 40.95 | 2.13 | 0.2407 | |
| BC | 102.96 | 1 | 102.96 | 5.35 | 0.1037 | |
| ABC | 73.81 | 1 | 73.81 | 3.84 | 0.1451 | |
| **Residual** | 57.73 | 3 | 19.24 | | | |
| Lack of Fit | 0.4667 | 1 | 0.4667 | 0.0163 | 0.9101 | not significant |
| Pure Error | 57.26 | 2 | 28.63 | | | |
| **Cor Total** | 2410.05 | 10 | | | | |

**Table S2**. ANOVA test for factorial model of the PDI of nanoparticles.

| Source | Sum of Squares | df | Mean Square | F-value | p-value | |
|---|---|---|---|---|---|---|
| **Model** | 0.0142 | 5 | 0.0028 | 35.56 | 0.0021 | significant |
| A-MM | 0.0029 | 1 | 0.0029 | 35.75 | 0.0039 | |
| B-SPC | 0.0004 | 1 | 0.0004 | 5.09 | 0.0870 | |
| C-P68 | 0.0074 | 1 | 0.0074 | 92.58 | 0.0007 | |
| AC | 0.0013 | 1 | 0.0013 | 16.63 | 0.0151 | |
| BC | 0.0022 | 1 | 0.0022 | 27.73 | 0.0062 | |
| Curvature | 0.0026 | 1 | 0.0026 | 32.02 | 0.0048 | |
| **Residual** | 0.0003 | 4 | 0.0001 | | | |
| Lack of Fit | 0.0002 | 2 | 0.0001 | 1.93 | 0.3407 | not significant |
| Pure Error | 0.0001 | 2 | 0.0001 | | | |
| **Cor Total** | 0.0170 | 10 | | | | |

**Table S3**. ANOVA test for factorial model of the Zeta potential of nanoparticles.

| Source | Sum of Squares | df | Mean Square | F-value | p-value | |
|---|---|---|---|---|---|---|
| **Model** | 180.33 | 5 | 36.07 | 28.38 | 0.0011 | significant |
| B-SPC | 129.20 | 1 | 129.20 | 101.67 | 0.0002 | |
| C-P68 | 6.35 | 1 | 6.35 | 5.00 | 0.0756 | |
| AC | 2.39 | 1 | 2.39 | 1.88 | 0.2288 | |
| BC | 21.35 | 1 | 21.35 | 16.80 | 0.0094 | |
| ABC | 21.03 | 1 | 21.03 | 16.55 | 0.0097 | |
| **Residual** | 6.35 | 5 | 1.27 | | | |
| Lack of Fit | 2.96 | 3 | 0.9875 | 0.5824 | 0.6816 | not significant |
| Pure Error | 3.39 | 2 | 1.70 | | | |
| **Cor Total** | 186.68 | 10 | | | | |

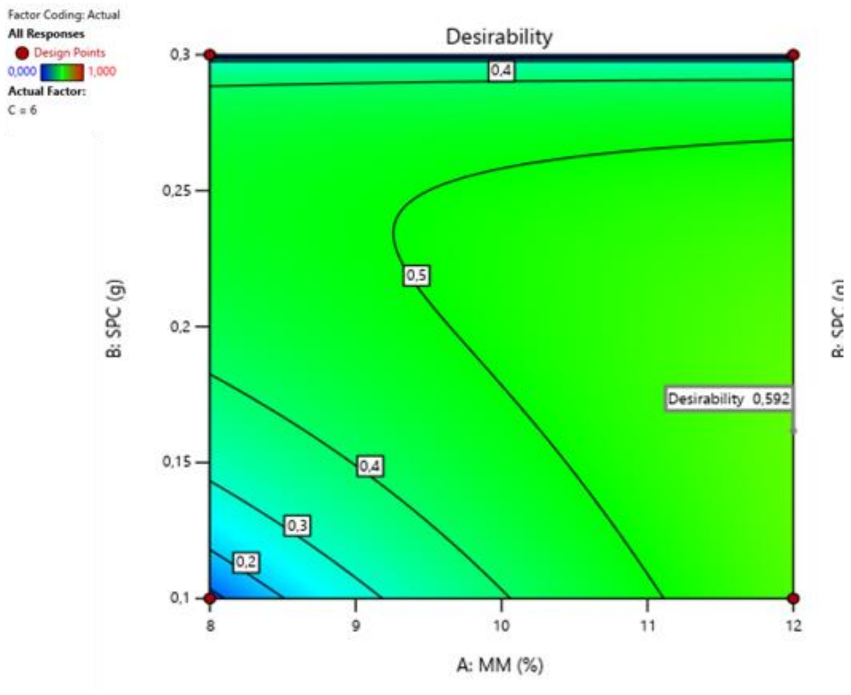


**Figure S2 –** Desirability graph of the NLC-CBD-PTX factorial design.
